# Microscopical and molecular characterization of the infection cycle of *Phytophthora betacei* during disease development on tree tomato (*Solanum betaceum)*

**DOI:** 10.1101/2025.09.11.675658

**Authors:** Natalia Guayazán Palacios, Juliana González Tobon, Maria Camila Buitrago, Daniel Bautista, Laura Natalia Gonzalez-Garcia, Martha Cárdenas Toquica, Maria Fernanda Mideros, Sebastian Schornack, Silvia Restrepo

**Author notes:** **Corresponding author: Silvia Restrepo**, Boyce Thompson Institute, 533 Tower Rd, Ithaca, NY. These authors contributed equally to this work and are considered co-first authors (Natalia Guayazán Palacios, Juliana González Tobon).

## Abstract

*Phytophthora betacei* is a recently described oomycete plant pathogen closely related to *Phytophthora infestans sensu stricto*. This plant pathogen naturally infects tree tomato (*Solanum betaceum*) but has not been reported on tomatoes and potatoes, the primary hosts of *P. infestans*. The aim of this study was to characterize the infection cycle of *P. betacei* using microscopy and molecular approaches. Several strains were inoculated in susceptible tree tomato plants and disease progression was monitored via six epidemiological parameters. Although different *P. betacei* strains displayed a highly variable disease phenotype, the most aggressive one was chosen for further plant inoculations. Samples at different time points of the infection cycle were analyzed at the cellular level via light and scanning electron microscopy (SEM) and at the molecular level via qRT-PCR of infection-stage-specific markers. The infection cycle of *P. betacei* differed from that of *P. infestans* in having a longer biotrophic stage, larger lesions, and higher sporulation capacity. Additionally, *P. betacei* transcriptomic profiles were monitored along the infection cycle via RNAseq and evidenced a changing expression landscape that supports an elongated hemibiotrophic transition and a clear distinction from what is being expressed in the mycelium or the sporangia. This study provides novel insights into the interaction between *P. betacei* and *S. betaceum*.

## Introduction

Remarkable efforts have been made in recent years to understand the mechanisms underlying pathogenesis and host colonization during plant disease. In terms of host colonization, only the pathogens that are able to establish and evade recognition, or suppress host defense mechanisms, are successful (Staskawicz 2001). To accomplish this, plant pathogens use diverse strategies. For example, bacteria can only gain access to the plant through natural openings (stomata, hydathodes) or wounds. Then, they establish themselves in the apoplast. In contrast, fungi and oomycetes can actively penetrate the plant epidermal cells and extend hyphae to eventually invaginate feeding structures (haustoria) in the host cell plasma membrane (Hogenhout et al. 2009). During these colonization and establishment processes, intimate molecular communication between the host and the pathogen occurs, leading to either plant resistance or disease (Jones and Dangl 2006).

Within the oomycetes, important research has been conducted concerning host colonization and pathogenicity (Hardham 2001; Kamoun 2007); especially for species within the *Phytophthora* genus (Bozkurt et al. 2012; Fawke et al. 2015; Kamoun 2007; Morgan and Kamoun 2007). This genus is considered one of the most devastating plant pathogens (Kamoun 2006) and comprises more than 100 species that have received special attention due to their economic and ecological impact (Martin et al. 2012). Some of the most representative species—and the diseases they cause—include: *P. infestans* (potato and tomato late blight), *P. capsici* (blight and fruit rot of pepper)*, P. sojae* (stem and root rot of soybean) (Lamour et al. 2007), *P. ramorum* (sudden oak death) (Grünwald et al. 2012), and *P. palmivora* (root rot of several plants, including oil palm) (Evangelisti et al. 2017).

Early studies of disease establishment by *Phytophthora* pathogens mapped out the major events that occur during the infection of host plants (Hardham 2001). As a whole, they showed that all *Phytophthora* species are hemibiotroph pathogens (Agrios 2005; Fawke et al. 2015). This means that they start the infection by feeding on live plant tissue (biotrophy) and then transition to killing the plant and feeding on its dead tissue (necrotrophy) later in the cycle. However, more recent research has expanded our understanding of hemibiotrophic infections via cell and molecular biology approaches (cf. Avrova et al. 2003; Avrova et al. 2008; Chen et al. 2014; Chen et al. 2013; Hardham 2001; Hayden et al. 2014; Judelson et al. 2008; Jupe et al. 2013; Kunjeti et al. 2012; Le Fevre et al. 2016; Potato Genome Sequencing Consortium. 2011; Ye et al. 2011). This revealed important differences among *Phytophthora* species regarding timing of each infection stage and the formation of stage-specific structures.

Recently, phylogenetic analyses, population genetics, and morphological approaches allowed the description of a new species: *Phytophthora betacei* (Mideros et al. 2018). This species coexists with *P. infestans* in South America and has been associated with late blight on tree tomato (*Solanum betaceum*), a semi-domesticated South American fruit crop. Interestingly, *P. betacei* has not been reported infecting potatoes or tomatoes (the main hosts of *P. infestans,* its sister species), suggesting that this species originated through ecological speciation by host specialization (Mideros et al. 2018; Restrepo et al. 2014). The transcriptional landscape of *S. betaceum* during the *P. betacei* infection was explored by Bautista et al. (2021) and showed how the plant switches its activated genes in response to the stage of infection and therefore, the attack mechanisms of the pathogen. However, the transcriptional response of the pathogen has not been yet explored.

This study aimed to characterize the infection cycle of *P. betacei* in its natural host, *S. betaceum*. For this, we first described the infection via multiple epidemiological parameters. Then, we thoroughly described the progression of the disease at a cellular level (via light and scanning electron microscopy) and at a molecular level (via the qRT-PCR of two infection-stage-specific markers). Lastly, we assessed the transcriptomic profiles of the pathogen at different points of the infection cycle.

## Materials and Methods

### Strain selection and maintenance

Ten *P. betacei* strains were selected from the *Phytophthora* collection maintained in the Natural Museum at Universidad de Los Andes (Supplementary Table 2). Those strains were maintained in tree tomato agar medium (1.8% bacteriological agar, 1.8% sucrose, 0.05% CaCO_3_, 20% tree tomato juice) at 18 °C. Three *P. infestans* strains isolated from potato fields (Supplementary Table 2) were included to compare infection cycles between species and were maintained in potato dextrose agar (PDA) (Oxoid, Waltham, MA, USA) and clarified V8 agar (2% bacteriological agar, 10% clarified V8, 0.1% CaCO_3_) at 18 °C.

### Growth conditions of plant material

Susceptible tree tomato (*S. betaceum*) plants of the ‘Comun’ variety were used for all experiments and grown from certified seeds (Impulsemillas, Bogotá, Colombia). All seeds were submerged in distilled water for 24 hours before germination according to the manufacturer. Seeds were germinated in peat and then transplanted to individual pots with potting soil. Germination and growth were conducted under greenhouse conditions (12 h/12h light period, 18-25 °C). All experiments in this study were performed on eight-to ten-week-old tree tomato plants.

### Plant inoculation assays

To ensure strain virulence, all *P. betacei* and *P. infestans* strains were transferred from *in vitro* cultures to leaves (‘Comun’ variety for tree tomato and ‘Suprema and Capiro’ varieties for potato, respectively) before using them for experimental inoculation assays. This was done via four 20 µL droplets of a non-adjusted sporangial suspension coming from the *in vitro* cultures and placed on the abaxial side of the leaves. The inoculated leaves were then placed inside petri dishes with a humid paper towel, creating moist chambers that were then incubated at 18°C until sporulation was observed.

Sporangial suspensions were prepared from sporangia growing on leaves using 1 mL of distilled water and adjusted to concentrations of 3 x 10^5^ sporangia mL^-1^ using a hemocytometer. The sporangial solutions were then incubated at 4°C for 3h to promote zoospore release (Aylor et al. 2007). Three leaves per plant were carefully drop-inoculated on their abaxial side using four 20 µL droplets of the suspensions. Three plants were inoculated per strain, as technical replicates, for a total of nine inoculated leaves. Inoculated plants were then kept inside a growth chamber (Percival, Perry, Iowa) at 17 ± 2 °C, 80% relative humidity and 12 h/12 h light period. Plants were individually covered with a clear plastic sheet throughout the entire experiment to maintain high humidity and prevent cross-contamination. The experiment was performed three times for a total of nine inoculated plants per strain.

### Epidemiological parameters of the infection cycle

Progression of disease caused by each strain was monitored for nine days using six epidemiological parameters: lesion area (LA, mm^2^ at day nine), lesion growth rate (LGR, mm^2^ day^-1^), incubation period (IP, hours post inoculation or hpi), sporulation capacity (SC, sporangia mm^-2^), latency period (LP, hpi), and sporulation rate (SR, sporangia mm^-2^ day^-1^) (Mideros et al., 2018).

Specifically, LA was determined using ImageJ 1.49 software (Schneider et al. 2012) to measure the necrotized leaf area. LGR was estimated by regressing LA for each lesion on each plant over time. The slope of the regression line was used as a proxy of LGR. IP was the time between inoculation and the first appearance of symptoms. SC was determined by rinsing sporangia from sporulating lesions into 1 mL of distilled water, counting them with a hemocytometer, and then dividing that number of sporangia over lesion area. LP was the time between inoculation and first sporulation. Lastly, SR was calculated by dividing sporulation by the time elapsed since the inoculation.

### Statistical analyses

Summary statistics were calculated for each epidemiological parameter per strain and per species. Subsequently, the normality of the residuals was assessed using the function “shapiro.test” available in the package ‘stats’ in R (R Core Team 2021). This test indicated that the data was not normally distributed (p < 0.01) and thus a Kruskal-Wallis test was performed to identify significant differences among strains and between species. This was followed by a Nemeyi post hoc test from the Pairwise Multiple Comparison of Mean Ranks Package (‘pmcmr’) (Pohlert 2014) to assess particular differences among strains. All p*-*values were corrected for multiple comparisons using the Bonferroni method and all the analyses were performed on pooled data from independent experiments with similar results.

### Clustering

We established whether the strains from *P. betacei* and *P. infestans* behaved biologically similar based on their epidemiological components. To this end, we visualized them all on a bidimensional plane and estimated the correlations using Spearman’s correlation coefficients. A Linear Discriminant Analysis (LDA) was implemented using a matrix of six epidemiological variables that did not exhibit collinearity. The analyses were conducted using the “lda” function from the package ‘mass’ in R (Venables and Ripley, 2002).

Lastly, to correctly identify groups of strains showing similar behaviour, a K-means analysis was implemented over the values obtained from the LDA (MacKay 2003). First, the number of ideal clusters *K* for the data was selected by implementing the elbow method, which relies on the explained variance. Here, the variance was measured as the sum of squares within a cluster and the maximum number of clusters allowed was 14, assuming each strain would group into a unique cluster. Once the *K* value was selected, the “kmeans” function from the package ‘stats’ in R (parameters set.seed = 30, *K*=5, nstart = 50) was used to assign data points to clusters. One *P. betacei* and one *P. infestans* strain belonging to highly different clusters were selected for further study.

### Light and scanning electron microscopy

To assess the infection cycles at the cellular level, two sets of leaf discs were excised at multiple time points (3, 6, 9, 12, 24, 48, 72, and 96 hpi) and used for light and scanning electron microscopy.

For light microscopy, one set of discs was subjected to trypan blue staining. Briefly, the discs were fixed in acetic acid (99%) / ethanol (96%) (3:1) until they had been fully decolorized (approximately 24 h, changing the decolorizing solution every 12 h), followed by dehydration in an ethanol series (50% and 70%) for 24 h each. Staining was then performed using 0.04% trypan blue in lactophenol/ethanol (96%) (1:2) solution for 2 minutes. They were then vacuum infiltrated at room temperature for 5 min and incubated in the trypan blue solution for another 2h at 4°C. Leaf pieces were mounted in glycerol 50% and visualized using a light microscope (ZEISS Axioskop 40, Goettingen, Germany).

For scanning electron microscopy, discs were fixed with 2.5% glutaraldehyde in 0.1 M Sorenson’s buffer (pH= 7.2) for 24 h at room temperature. Subsequently, they were washed twice, each time for 10 minutes, in the same buffer. After fixation, the samples were dehydrated in a graded ethanol series as follows: 50%, 70%, 90%, and 95% for a period of 30 min in each series, and two changes of 2h each in a 100% series. The samples were critical-point dried using CO_2_. The fixed material was mounted on stubs using double-sided carbon tape and coated with gold/palladium in a sputter coater system in a high-vacuum chamber for 150 s at 9 mA (Soylu, Soylu and Kurt 2006). The samples were then visualized using a Scanning Electron Microscope (SEM) accelerating voltage 20kv (JEOL Microscope, model JSM 6490-LV, Peabod, MA, USA).

### Total RNA extractions

Three pieces of leaf samples were also excised at multiple time points (6, 12, 18, 24, 72, and 96 hpi), pooled, and placed in liquid nitrogen immediately. Tissue from uninoculated leaves, mycelia, and sporangia were also harvested. The RNase Plant MiniKit® (Qiagen, Germantown, MD, USA) was used to extract total RNA from all these samples according to the manufacturer’s instructions. Purified RNA was treated with DNase I and its integrity and yield were measured using gel electrophoresis and a 2100 bioanalyzer (Agilent, Waldbronn, Germany).

### Quantitative RT-PCR

To assess the infection cycles at the molecular level, total RNA from inoculated leaves was then used for qRT-PCR to assess expression patterns of infection-stage specific markers. cDNA was generated using the High-Capacity cDNA Reverse Transcription Kit (Applied Biosystems, Foster City, CA, USA) with 1 µg o]RNA. A qRT-PCR was performed to examine the gene expression levels of the haustorium-specific membrane protein *Hmp1* (Avrova et al. 2008) and the cell cycle regulator *Cdc14* (Ah Fong and Judelson 2003) by using Maxima SYBR Green qPCR (Thermo Scientific, Waltham, MA, USA) and a 7500fast thermocycler (Applied Biosystems, Foster City, CA, USA).

These marker genes were first identified on the *P. betacei* genome (Ayala-Usma et al. 2021) using Basic Local Alignment Search Tool BLAST (Altschul et al., 1990) with the homologues found in *P. infestans* as a query. The Actin gene *PbactA* was used as an endogenous control gene (Supplementary Table 3) and mycelium was used as the calibrator. Reactions consisted of a final volume of 15 µL including 2 µL cDNA 0.45 µL of each 0.3 µM primer, 5 µL of Maxima SYBR green qPCR master mix without ROX, 0.03 µL of 10 nM ROX, and nuclease-free water to complete the volume. The PCR parameters were as follows: 95 °C, 15 min; 95 °C, 15 s; 60 °C, 30 s; 72 °C, 30 s (40 cycles). Melting curve analyses were performed on every run to confirm a single product (60–95°C) reading every 1°C (Bos et al. 2010). Quantification analyses were performed according to the Pfaffl method (Pfaffl 2001) and using the software REST (Pfaffl et al. 2002).

### Library preparation and RNA sequencing

To assess the transcriptomic profile of *P. betacei* along its infection cycle, total RNA from inoculated leaves, uninoculated leaves, mycelium, and sporangia was also used for sequencing pathogen transcripts as disease progressed. Library preparation and sequencing were performed by Novogene Inc. Briefly, *P. betacei* messenger RNAs (mRNAs) were purified using Poly(A) selection from the total RNA sample and then fragmented. Then, a cDNA library was prepared with the non-directional NEBNext® Ultra™ RNA Library Prep Kit for Illumina® (New England Biolabs, Hitchin, UK) according to the manufacturer’s protocol and sequenced on an Illumina HiSeq4000 system (Illumina, San Diego, CA) over a paired-end library (insert size of ∼250 -300 bp) for three biological replicates per time point.

### *de novo* transcriptome assembly and analysis for *P. betacei*

#### Read cleanup

Reads (∼1342 M) from all samples were quality-assessed using the FASTQC tool v.0.11.2 (Babraham Bioinformatics, Cambridge, UK). Then, potential contaminants were identified by screening the reads against the SILVA databases using ‘bbduk.sh’ from the ‘BBtools’ package (kmer = 27) (Joint Genome Institute). The sequences of identified contaminants were obtained from NCBI and used to further clean the reads using the same tool. Lastly, a step of trimming and filtering was performed using Trimmomatic v.0.36 (Bolger et al., 2017) with the following parameters: threads 5, phred33, HEADCROP:10, MINLEN:80. The remaining (∼1310 M) reads were subjected to FASTQC and used in downstream analyses.

#### Read filtering

Clean reads from the uninoculated plant material were used in a *de novo assembly* of a reference transcriptome for *S. betaceum* using Trinity v2.4 (Grabherr et al. 2011; Haas et al. 2013) (with kmer=25). The *S. betaceum* transcriptome assembly was then used to exclude plant reads from the inoculated samples’ libraries by implementing the pseudoalignment function ‘pseudo’ from Kallisto (Bray et al. 2016). Remaining reads were subject to a mapping round of HISAT2 (Kim et al. 2015) against the *S. tuberosum* (v4.03), *S. lycopersicum* (v3.0), and *Caspsicum annuum* (v2.0) reference genomes, given their proximity to *S. betaceum*.

#### Construction of the P. betacei de novo transcriptome

Cleaned reads from the axenic tissues (mycelium and sporangia) were then used in a *de-novo* assembly of the *P. betacei* transcriptome with the same parameters as the *S. betaceum* assembly. *In silico* read normalization was used during transcriptome assembly due to the large number of input reads.

#### De novo assembly statistics and transcriptome completeness

General statistics of the assembly were obtained using the native “TrinityStats.pl” script and independently using the software Transrate (Smith-Unna et al. 2016). Furthermore, the number of full-length transcripts was estimated by examining the extent of top-matching BLASTX (Altschul et al. 1990) alignments against the SwissProt database, the percent of the target being aligned to the best matching Trinity transcript, and by grouping blast hits to improve sequence coverage (Haas et al. 2013). These scripts are available with the Trinity utilities. In addition, the completeness of each assembly was assessed with BUSCO (Simão et al. 2015) according to the presence of single copy orthologs from Eukaryota.

#### P. betacei transcriptome annotation

Predicted transcripts were mapped to a previously reported *P. betacei* genome (Ayala-Usma et al. 2021) using BLASTN (Altschul et al. 1990). Matches were filtered by 75% identity, 1e-5 E-value, and 50% query coverage. The longest transcript isoform and the best match per gene were retained. Functional annotation, including Gene Ontology terms, was extracted from a previous complete genome annotation (Ayala-Usma et al. 2021).

#### Assessment of transcript abundance and differential expression analysis

To identify differentially expressed genes, we first quantified pathogen transcripts by quasi-mapping the reads from each sample against the *P. betacei de novo* transcriptome using Salmon (Patro et al. 2017). Quantification was expressed as Transcripts Per Million (TPM) and further aggregated to the gene level using the ‘tximport’ package in R (Soneson et al. 2015) for gene-level differential expression analysis with DESeq2 (Love et al. 2014). Genes with less than five mapped reads were removed. Then, differentially expressed genes (DEGs) were calculated at each time point when compared to the mycelium, filtered by a p-value ≤ 0.05 and a log2FoldChange (log2FC) ≥ |*2*|, and normalized using the RLD (regularized log transformation) method (Love et al. 2014). A Principal Components Analysis was performed using the top500 differentially expressed genes and the *plotPCA* function of DESeq2. A heatmap of this same set of genes was built using the package ‘pHeatmap’ in R (Kolde 2015) and upset plots of the number of significantly up and downregulated genes were built using the package ‘upsetR’ in R (Conway 2017).

#### Analysis of P. betacei effector expression along the infection cycle

To identify the expression profiles of *P. betacei* effectors along the infection cycle, we first extracted the sequences for the 979 effectors previously reported for *P. betacei* (Ayala-Usma et al. 2021; Rojas-Estevez et al. 2020). Among them, 793 were annotated as RxLR and 144 were annotated as Crinkler (CRN) (Ayala-Usma et al. 2021). These effector sequences were used as queries and were blasted, as done previously, against our *de novo* transcriptome to identify the transcript isoforms that matched them. Afterwards, RNAseq counts were filtered for these genes to analyze the expression profiles of RxLR and CRN effectors across the infection cycle.

#### Validation of transcriptomic profiles via qRT-PCR

The expression of selected differentially expressed genes was validated by qRT-PCR as indicated above. In this case, five genes were evaluated (Supplementary Table 4). These genes were chosen because of their diverse roles in the infection cycle of *P. infestans*. In this case, the elongation factor 1 gene (*EF-1)* was used as the endogenous control gene (Supplementary Table 4) and mycelium was used as the calibrator (Bos et al. 2010). Quantification analyses were performed as previously described.

## Results

### *P. betacei* isolates display a highly variable phenotype while colonizing a susceptible host

To understand how the disease on tree tomato progresses, we measured six epidemiological parameters on plants infected with *P. betacei* and *P. infestans* strains. Most strains from *P. betacei* displayed significant differences in their LA (p < 2.3 x 10^-6^), IP (p < 1.86 x 10^-10^), and SC (p < 1.94 x 10^-4^) when compared to each other and when compared to strains from *P. infestans.* In contrast, no differences were detected among *P. infestans* strains (Table 1). As a species, *P. betacei* differed significantly from *P. infestans* for LA (p < 4.655 x 10^-5^), IP (p < 1.577 x 10^-6^), SC (p < 2.2 x 10^-16^), and SC (p < 7.65 x 10^-5^) (Supplementary table 1).

**Table 1.** Median value of each evaluated epidemiological parameter between species. The first and third quartiles are indicated in parentheses as a dispersion measurement.

| Epidemiological parameter | Species |  |
| --- | --- | --- |
|  | *P. betacei*<br>*n*=11 | *P. infestans*<br>*n*=3 |
| **Lesion Area (mm<sup>2</sup>)** | 18.8<br>(15.24-42.36) | 64.45<br>(33.23-86.90) |
| **Lesion Growth Rate (mm<sup>2</sup>.h<sup>-1</sup>)** | 0.087<br>(0.07-0.19) | 0.29<br>(0.15-0.40) |
| **Incubation Period (hpi)** | 81.19<br>(53.33-112.0) | 46.59<br>(40.0-51.0) |
| **Sporulation Capacity (sp.mm<sup>-2</sup>)** | 405.1<br>(0-509.7) | 47.14<br>(0-69.53) |
| **Sporulation Rate (sp.h<sup>-1</sup>)** | 39.8<br>(0-100.6) | 11.57<br>(0-25.63) |
| **Latency Period (hpi)** | 185.1<br>(0-216) | 154.3<br>(0-216) |

A linear discriminant analysis (LDA) using 5 of the 6 variables (LGR was excluded due to significant correlation to LA pval<0.01) revealed that *P. betacei* has variable infection pattern as indicated by the spread phenotypes, compared to *P. infestans* where all the strains grouped together. This observation was further supported by a k-means clustering analysis that identified three groups of strains (Fig. 1A). The largest differences were driven in the LA and SC in the first LD function (LD1, 76.95%), and by IP in the second (LD2, 12.33%). Overall, *P. betacei* has a high intra-species variety for LA, ranging from small to large areas (Fig. 1B). Although IP was more consistent and values tended to be high, there was also some variation (Fig. 1B). In contrast, strains from *P. infestans* have a similar performance when compared to each other regarding LA and IP (Nemeyi Test p > 0.01). Based on these observations we selected N9035 from *P. betacei* as a highly aggressive strain on the “Comun” cultivar, and Z32 from *P. infestans* as a comparison for cellular, and molecular analyses.

**Fig. 1.**
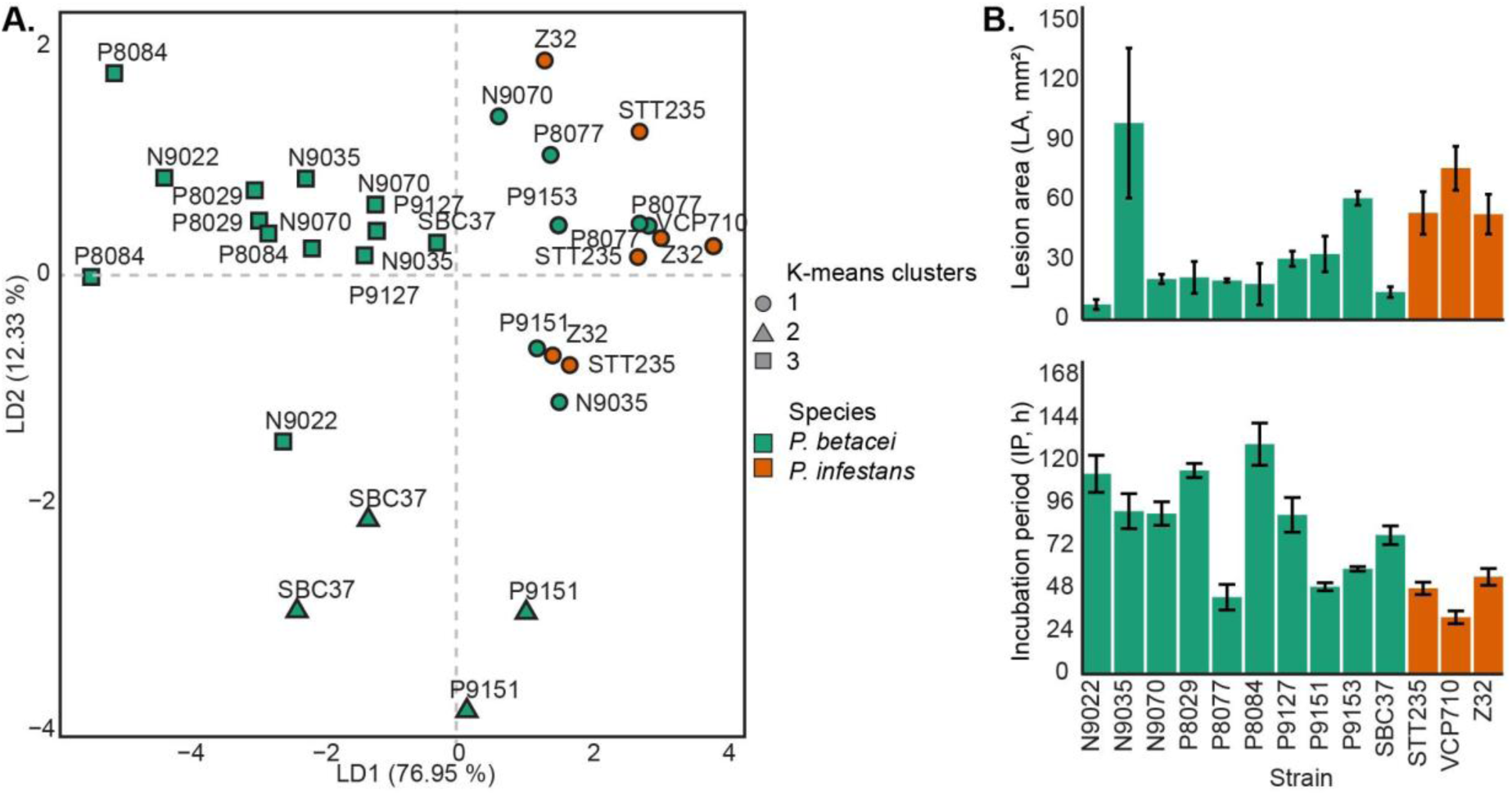
Clustering analysis of *Phytophthora* strains based on six epidemiological parameters. A) Linear Discriminant Analysis (LDA) showing the first and second linear discriminants. The *Phytophthora* strains were differentiated by Lesion Area (LA) and Sporulation Capacity (SC) on the first component (LD1=76.95%), and by Incubation Period (IP) on the second component (LD2=12.33%). Due to collinearity, Lesion Growth Rate (LGR) was excluded from the analysis. The linear discriminant values from each strain were used to assign 3 clusters. The ideal number of clusters was selected based on the total within the cluster sum of squares. Each cluster represents a group of similar strains. **B) Lesion area and incubation period.** Lesion area (mm^2^) and incubation period (hpi) from 10 *P. betacei* strains and 3 *P. infestans* strains. Error bars represent the standard error of the mean (SEM).

### *P. betacei* and *P. infestans* – tree tomato interactions feature a hemibiotrophic infection cycle

To understand how the *P. betacei -* tree tomato interaction looks at the cellular and molecular levels, as well as how it compares to the *P. infestans’* one, both microscopic and molecular analyses were performed.

N9035 and Z-32 presented differences in the timing of biotrophic and necrotrophic stages. In the early stages of infection (up to 72 hpi), *P. betacei* featured a biotrophic phase during which host tissue appeared healthy and unaffected (Figs. 2A-C), followed by a necrotrophic phase (>96 hpi) where water soaking, necrosis, and tissue collapse were evident (Fig. 2D-E). New sporangia were visible on the rim of the necrotic area at 120 hpi (Fig. 2E). Meanwhile, *P. infestans* rapidly switched from biotrophy to necrotrophy (>24 hpi) (Fig. 3A) and the tissue rapidly decayed (Fig. 3A-D). However, sporangia were not detected by direct visual inspection.

**Fig. 2.**
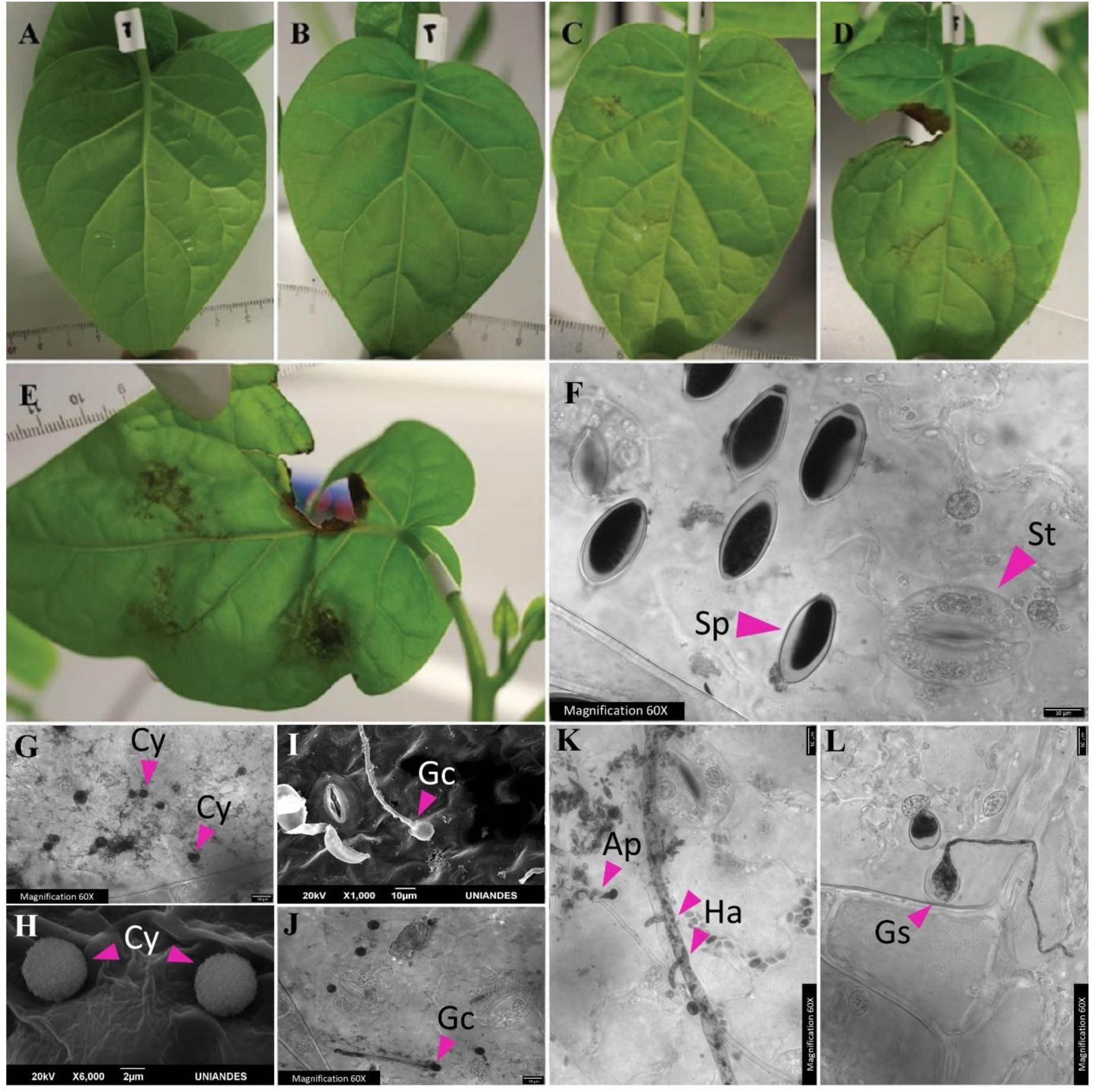
*Phytophthora betacei* features a hemibiotrophic lifestyle with an extended biotrophic phase A-E) Inoculated tree tomato plants after 24 (A), 48 (B), 72 (C), 96 (D) and 120 (E) hours post inoculation (hpi). **F-L)** Detail of the infection process of *P. betacei* under light (F, G, J, K, L) and Scanning Electron Microscopy (H, I) at 0 (F), 3 (G-H), 6 (I-J), 9 (K), 12 (L) hpi. Sp=Sporangia, St=Stomata, Cy=Cyst, Gc=Germinated Cyst, Ap=Appressorium, Ha=Haustoria, Gs= Germinated Sporangia. Microscopical images were trypan blue-stained.

**Fig. 3.**
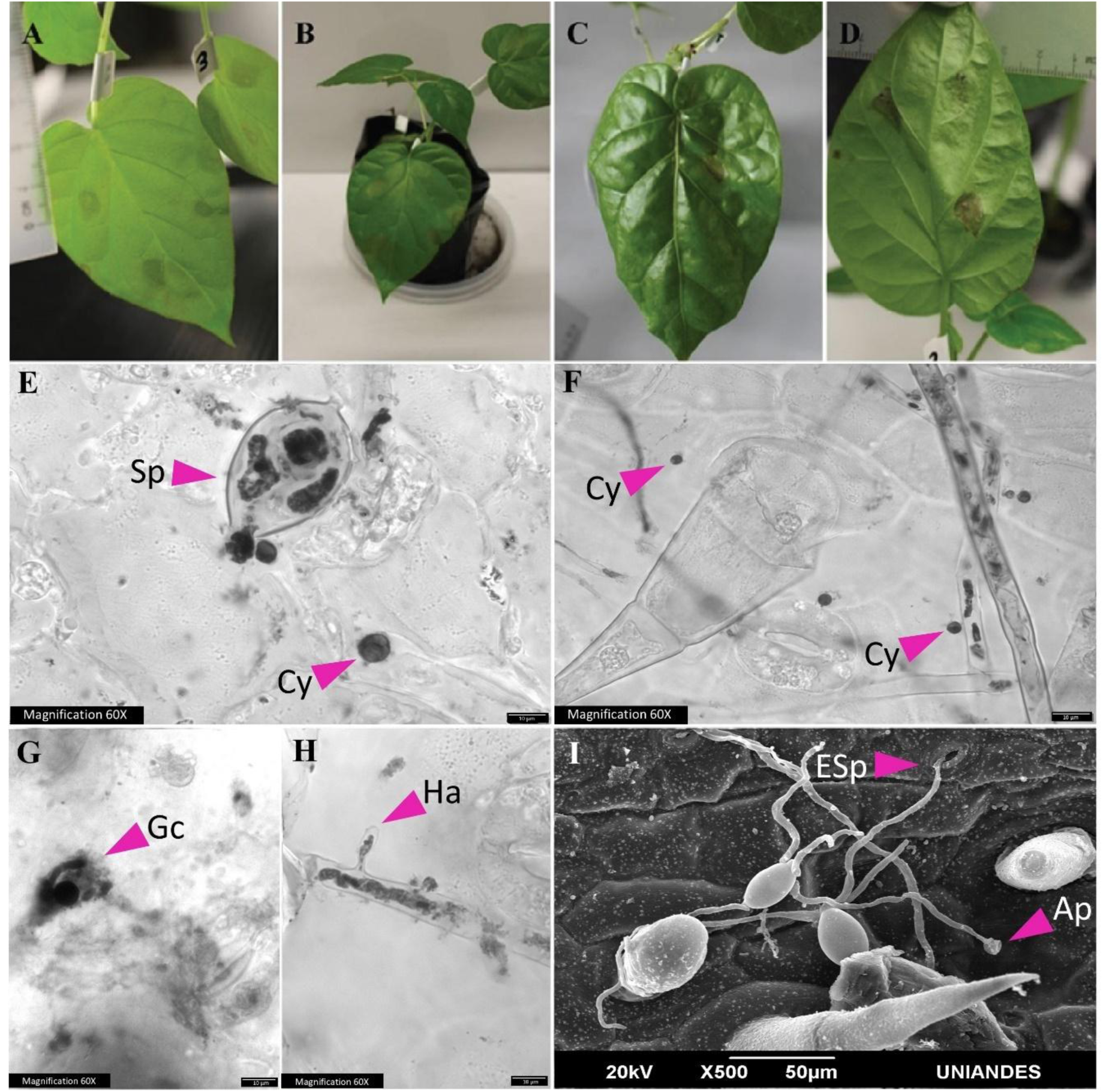
*Phytophthora infestans* features a hemibiotrophic lifestyle on a non-natural host. A-D) Inoculated tree tomato plants after 24 (A), 48 (B), 72 (C), 96 (D) hours post inoculation (hpi). **E-I)** Detail of the infection process of *P. infestans* under light (E-F) and Scanning Electron Microscopy (I) at 0 (E), 3 (F), 6 (G), 9 (H), 72 (I) hpi. Sp=Sporangium, St=Stomata, Cy=Cyst, Gc=Germinated Cyst, Ap=Appressorium, Ha=Haustorium Gs= Germinated Sporangia, Esp = Emerging sporangia.

When assessed microscopically, both strains were capable of forming appressoria, haustoria, and invasive hyphae. Cysts were observed as early as 3 hpi, germinated cysts at 6 hpi, and appressoria and haustoria >9 hpi (Figs. 2G-K and 3E-H). Microscopic inspection of leaf tissue in the later infection stages revealed significant colonization with the formation of sporangia 120 and 72 hpi for N9035 and Z-32, respectively (Figs. 2E and 3I). Interestingly, *P. betacei* did not appear to release the zoospores from the sporangium until after 3 hours of reaching the host, while *P. infestans* appeared to liberate the zoospores while on the sporangial solution previous to the inoculation (Figs. 2F and 3E).

Both direct and indirect germination of the sporangia led to the formation of appressoria and haustoria on asymptomatic tissue, suggesting a biotrophic interaction with the host. Later, tissue collapse and necrosis accompanied by sporulation indicated the necrotrophic phase and the end of the infection cycle. The longer time taken by *P. betacei* than by *P. infestans* to transition towards necrotrophy suggests a longer biotrophy period and, overall, a hemibiotrophic infection cycle.

### Varying expression of molecular markers supports differences in infection cycles of *P.* betacei and P. infestans on tree tomato

To determine if the timing of observed microscopic structures correlated with the timing of expression of marker genes, we measured the expression of the *P. betacei* homologs for Haustoria Membrane protein 1 (*Hmp1*) and Cell cycle phosphatase 14 (*Cdc14*). qRT-PCR results indicated that *Hmp1* was upregulated in both species at 12 hpi, but had a greater fold-change for *P. infestans* relative to mycelium. After this time, the gene became downregulated in *P. betacei* while peaking again for *P. infestans* at 72 hpi (Fig. 4A). On the other hand, *Cdc14* was highly induced later during the infection cycle for both species, peaking at 96 and 72 hpi, respectively (Fig. 4B). These results positively correlated with the disease progression and microscopic observations and suggested that the long biotrophic phase of *P. betacei* might be explained by the slower formation of early infection structures like cyst and haustoria.

**Fig. 4.**
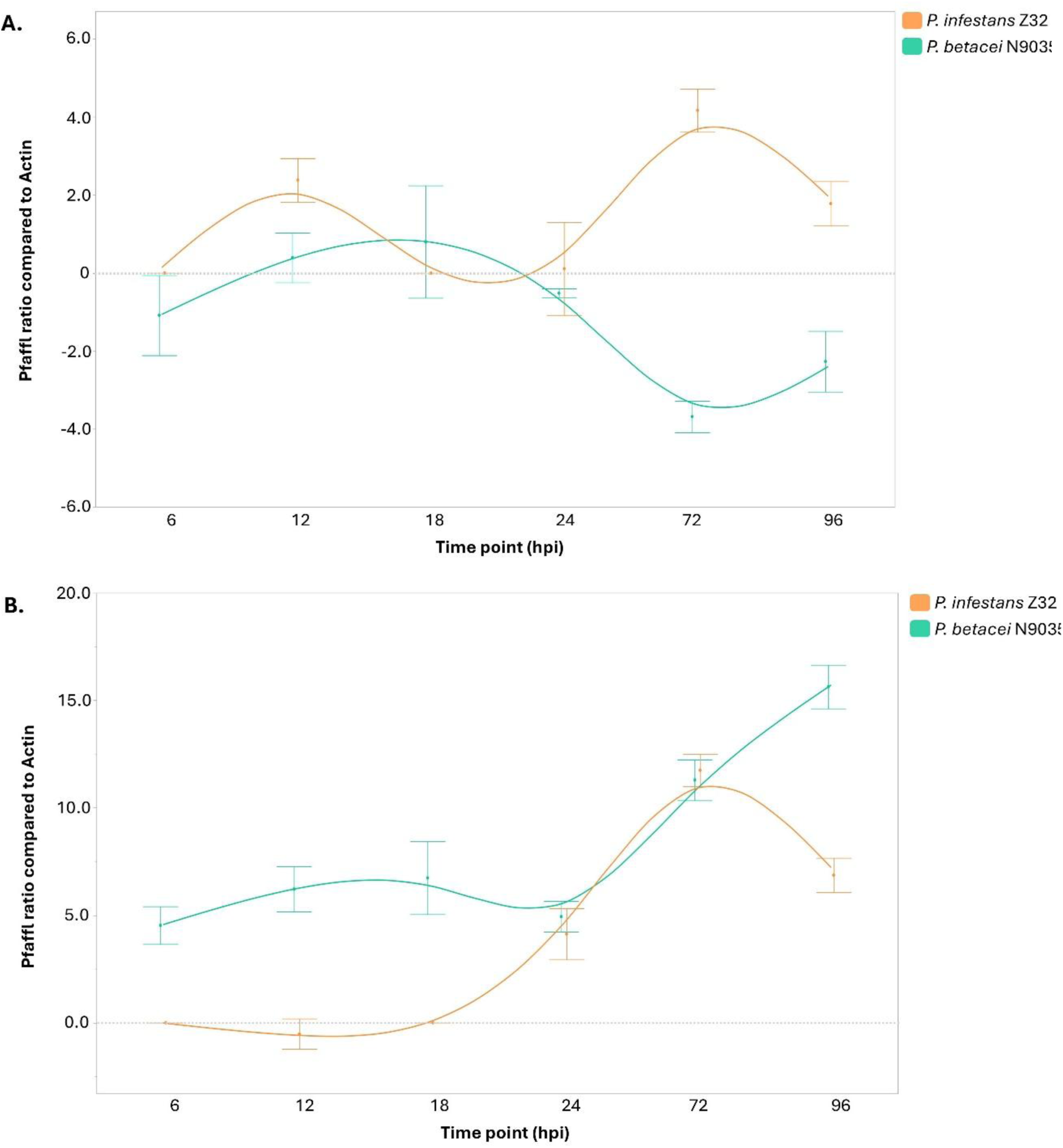
Monitoring of infection cycle stages by qRT-PCR. Expression relative to actin was assessed for the lifestyle marker genes Haustorium membrane protein *Hmp1* **(A)** and Cell cycle phosphatase *Cdc14* **(B)**, on time-course experiments for *P. betacei* (green) and *P. infestans* (orange). Dots represent the mean value of relative expression and bars represent the standard error (SE) across two independent repetitions of the experiment.

### The transcriptomic signature of P. betacei changes throughout its cycle on tree tomato

*P. betacei* gene expression was monitored along the infection cycle by creating a *de novo* transcriptome (Supplementary Files 1 and 2) of the pathogen and then aligning the reads from each sampled time point against it. A summary of the assembly statistics of the *de novo* transcriptomes can be found in Table 2. A principal component analysis (PCA) of the top500 DEGs revealed a major difference between the mycelium and sporangia samples when compared to all time points of inoculated leaves (Fig. 5A). By analyzing the inoculated plant samples, different groups became visible, separating those of the biotrophic stage (6, 12, 18, 24 hpi) from those of the necrotrophic stage (72 and 96 hpi) later in the cycle (Fig. 5B).

**Fig. 5.**
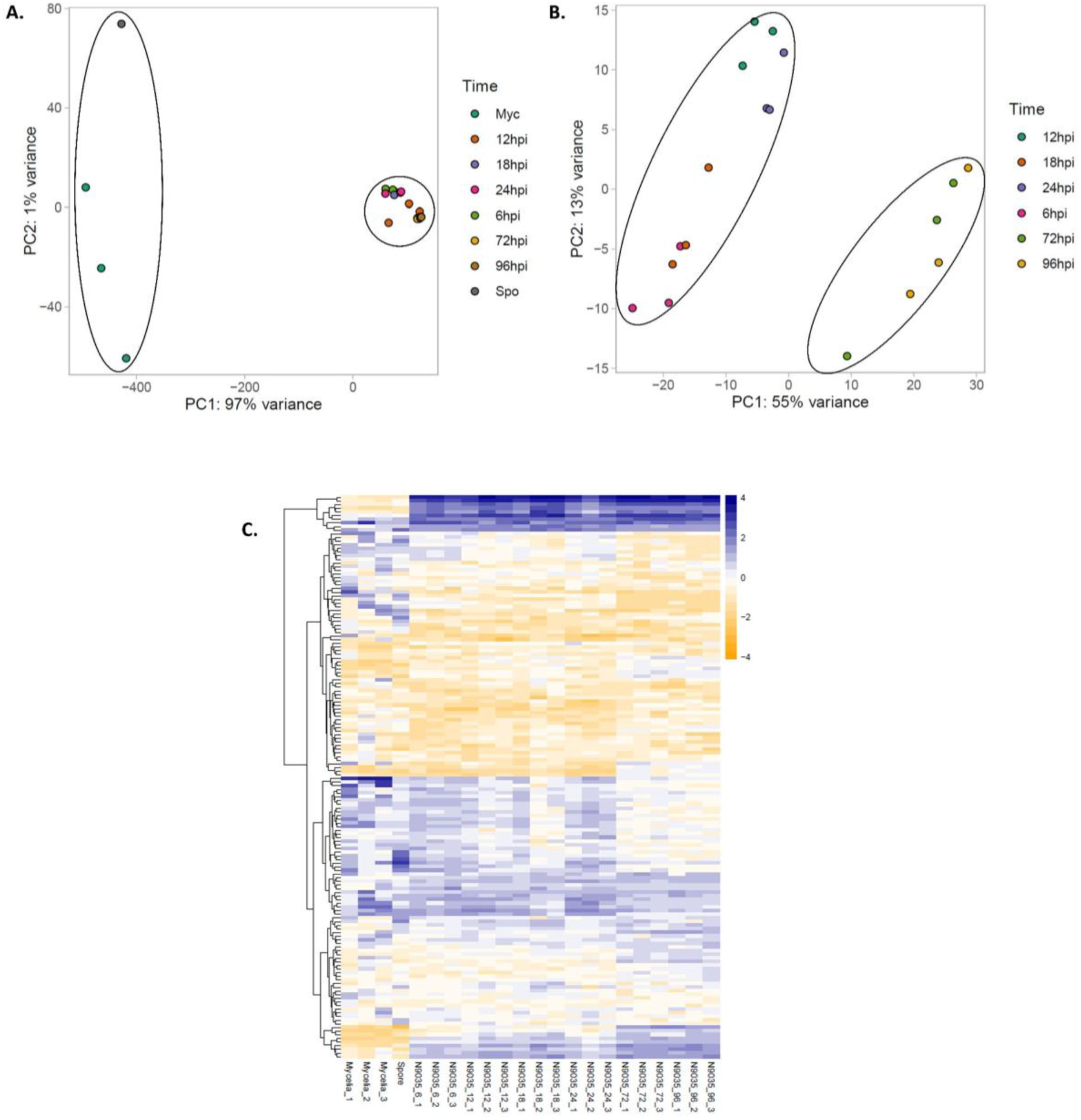
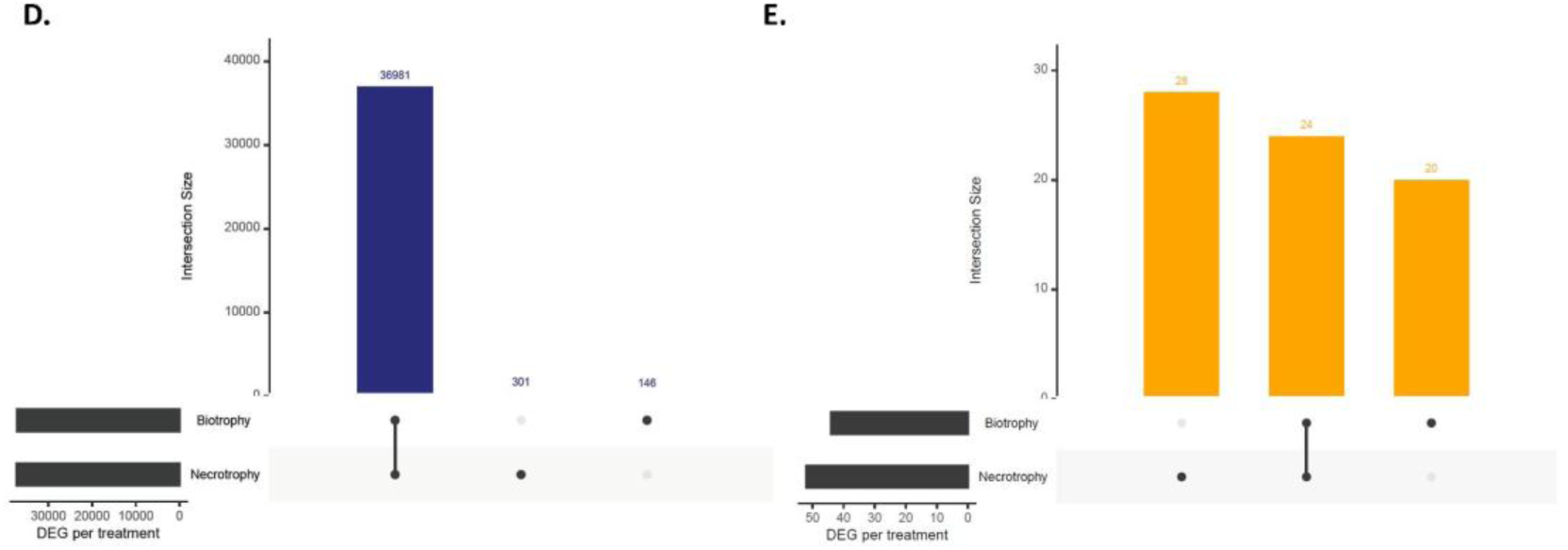
Transcriptomic analyses of *Phytophthora betacei* during tree tomato colonization. Principal components analysis (PCA) of the top500 differentially expressed genes (DEGs, host infection time point vs mycelium LFC ≥ 2, padj <0.05) across **(A)** all samples and **(B)** only samples from *P. betacei* infecting tree tomato (*S. betaceum*). **(C)** Heatmap of the mean-centered z-scores of the top500 DEGs of all *P. betacei* samples. UpSet plots indicate unique and shared **(D)** up-regulated and **(E)** down-regulated genes between biotrophy (6, 12, 18, 24 hpi) and necrotrophy (72 and 96 hpi), when compared to the mycelium.

**Table 2.** De novo transcriptome assembly statistics for *P. betacei* and *S. betaceum*.

| Metric | Value |  |
| --- | --- | --- |
|  | *P. betacei* axenic | *S. betaceum* leaf |
| Total Trinity genes | 53,079 | 113,756 |
| Total Trinity transcripts | 135,443 | 241,871 |
| Percent GC | 51.23 % | 38,81 % |
| Smallest contig | 201 | 201 |
| Largest contig | 43,434 | 14,798 |
| Statistics based on all transcripts |  |  |
| Contig N50 | 2,240 | 1691 |
| Median contig size | 702 | 541 |
| Average contig size | 1251,14 | 968,02 |
| Total assembled bases | 169,457,503 | 234,137,138 |
| Statistics based on the longest isoform per gene |  |  |
| Contig N50 | 1754 | 1247 |
| Median contig size | 408 | 389 |
| Average contig size | 885,47 | 728,42 |
| Total assembled bases | 4,699,844 | 82,862,089 |

Given how different the mycelium and sporangia were from the inoculated plant samples, we then generated a heat map of the top500 DEGs (Fig. 5C, Supplementary File 3). This heatmap shows that the transcriptomic profiles of the mycelium and the sporangia are very similar to each other, while very different from all the other samples. The inoculated samples, on the other hand, had similarities across them all (e.g., a clear group of upregulated genes represented in the upper blue panel) but differed enough between biotrophy (6, 12, 18, and 24 hpi) and necrotrophy (72 and 96 hpi) (Fig. 5C).

Overall, our DEGs included many uncharacterized proteins with diverse expression profiles (up/down regulated at different points of the cycle) (Supplementary File 4). This prevented the enrichment of GO terms in our results. However, examples of genes upregulated in the inoculated samples but downregulated in the mycelium and sporangia include a glucose transporter (PBET_0001694-RA) and a secreted protein (PBET_00016520-RA). While examples of genes downregulated in the inoculated samples but upregulated in the mycelium and sporangia include a cell division protein kinase (PBET_00003633-RA) and a Hsp70-like protein (PBET_00004705-RA).

Overall, 146 genes were upregulated exclusively in biotrophy, and 301 exclusively in necrotrophy, when compared to the mycelium (Fig. 5D). On the other hand, 20 genes were downregulated exclusively in biotrophy and 28 exclusively in necrotrophy, when compared to the mycelium (Fig. 5E). Although many of these genes were also uncharacterized proteins, three genes of interest were identified as differentially expressed (Fig. 6). Among them, the glucanase inhibitor protein 1 (*GIP*) was upregulated very early in the biotrophy (6 hpi), decreased in expression as this phase continued (between 12 and 18 hpi) and then increased its expression again at 24 hpi, which was maintained into the necrotrophic stage. The suppressor of necrosis 1 (*SNE1*) was clearly upregulated in the necrotrophic phase, with the peak starting at around 24 hpi. Lastly, the mediator of RNA Pol II subunit increased its expression progressively during the biotrophic phase and peaked towards the end of the cycle (72 and 96 hpi). The expression profiles of these genes across the cycle were also validated using qRT-PCR (Fig. 6).

**Fig. 6.**
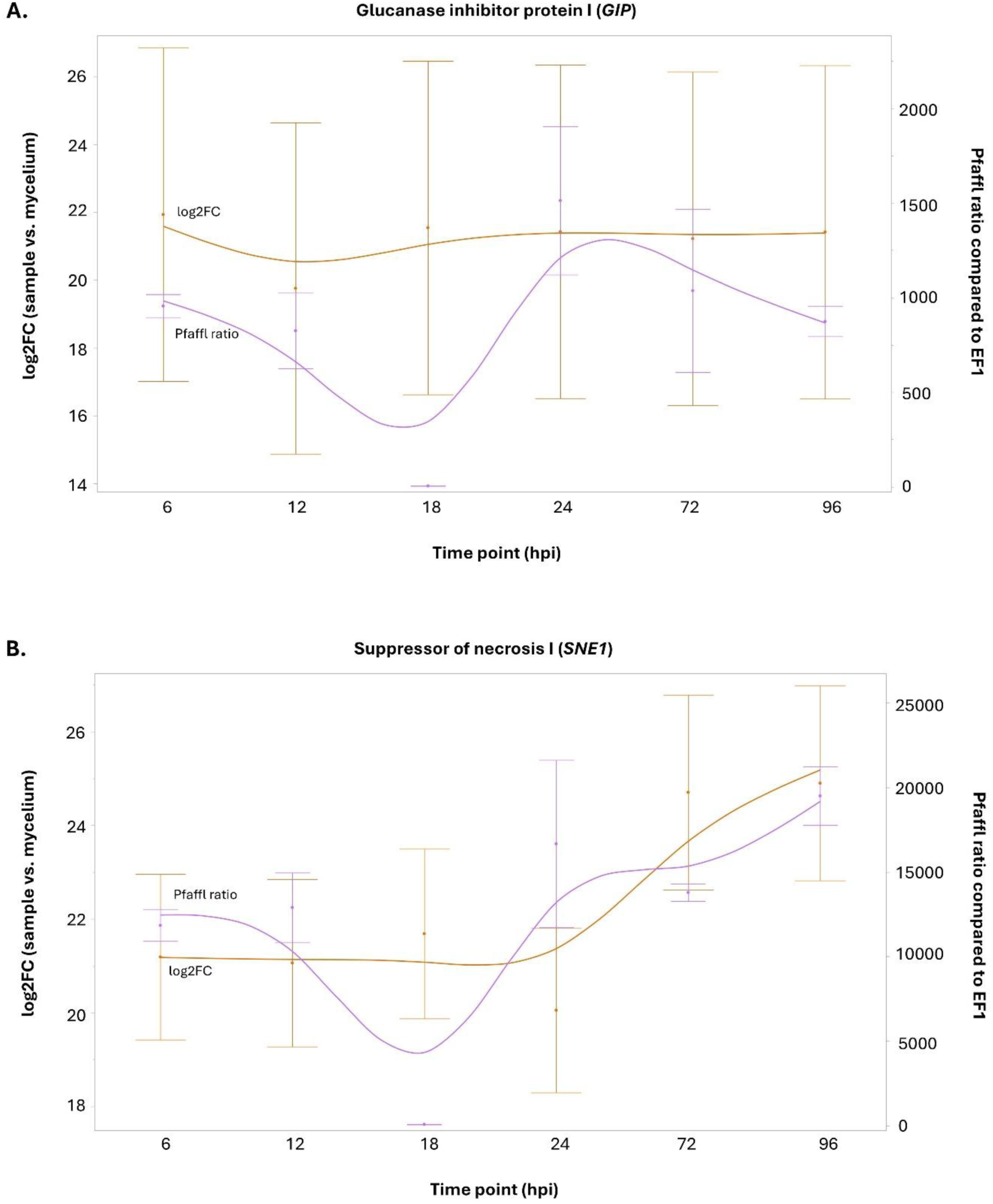
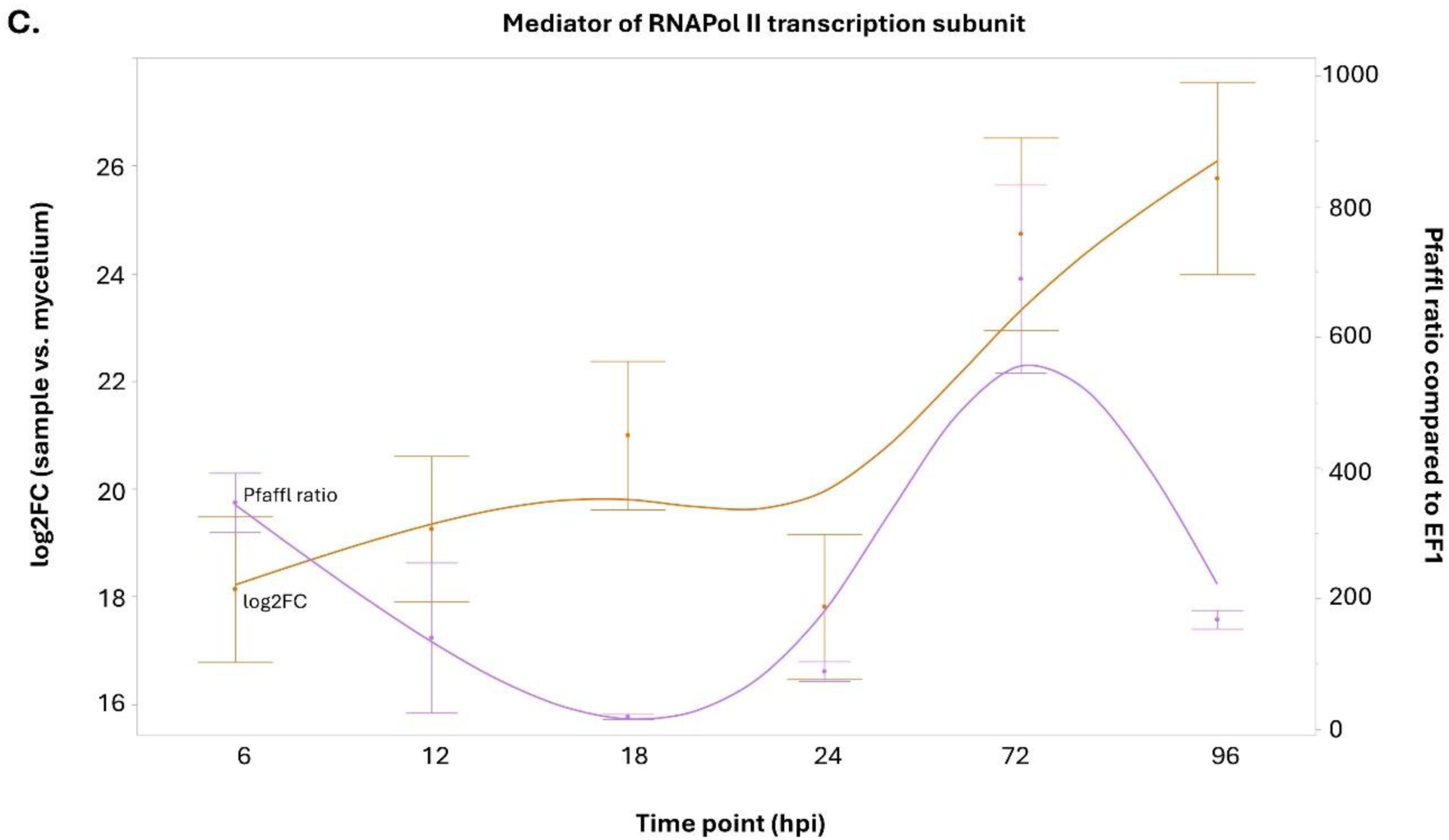
Validation of the transcriptomic profile of three genes of interest via qRT-PCR. Expression relative to the elongation factor 1 (EF-1) was assessed for the glucanase inhibitor protein 1 (GIP) **(A)**, the suppressor of necrosis I (SNE1) **(B)**, and the mediator of RNA Pol II **(C)**. Data points are either the Pfaffl ratio values (purple) or the log2fold change values (orange) of each gene at each time point when compared to the expression of EF-1, or its expression in the mycelium, respectively. Therefore, the purple line shows the pattern of expression in the qRT-PCR, and the orange line shows the pattern of expression in the RNAseq. Each data point is an average, bars represent the standard error (SE) across technical replicates (qRT-PCR) and biological replicates (RNAseq).

In our transcriptome, we identified 698 effector genes (Supplementary File 5) out of the 979 that had been reported previously (Ayala-Usma et al. 2021; Rojas-Estevez et al. 2020). Among them, 479 were RxLRs, 195 were CRNs and 24 were annotated as effectors but from no particular group. RxLRs and CRNs showed an overall trend of upregulation at 72 and 96 hpi, in comparison to the mycelium and early stages of the infection (˂72 hpi) (Figs. 7A and B). However, within RxLRs we identified more variability among members, having some that kept their expression constant during biotrophy and even the hemibiotrophic transition but peaked at 72hpi (Fig. 7C, PBET_00013572-RA), others upregulated at 72 and 96hpi (Fig. 7C, PBET_00019987-RA and PBET_00009358-RA), and others that peaked earlier in the infection cycle, at 24hpi (Fig. 7C, PBET_00019978-RA). On the other hand, CRNs showed a clearer trend where most of them had a semi-constant expression during biotrophy and hemibiotrophy, but strongly upregulated at 72 and 96 hpi (Fig. 7D).

**Fig. 7.**
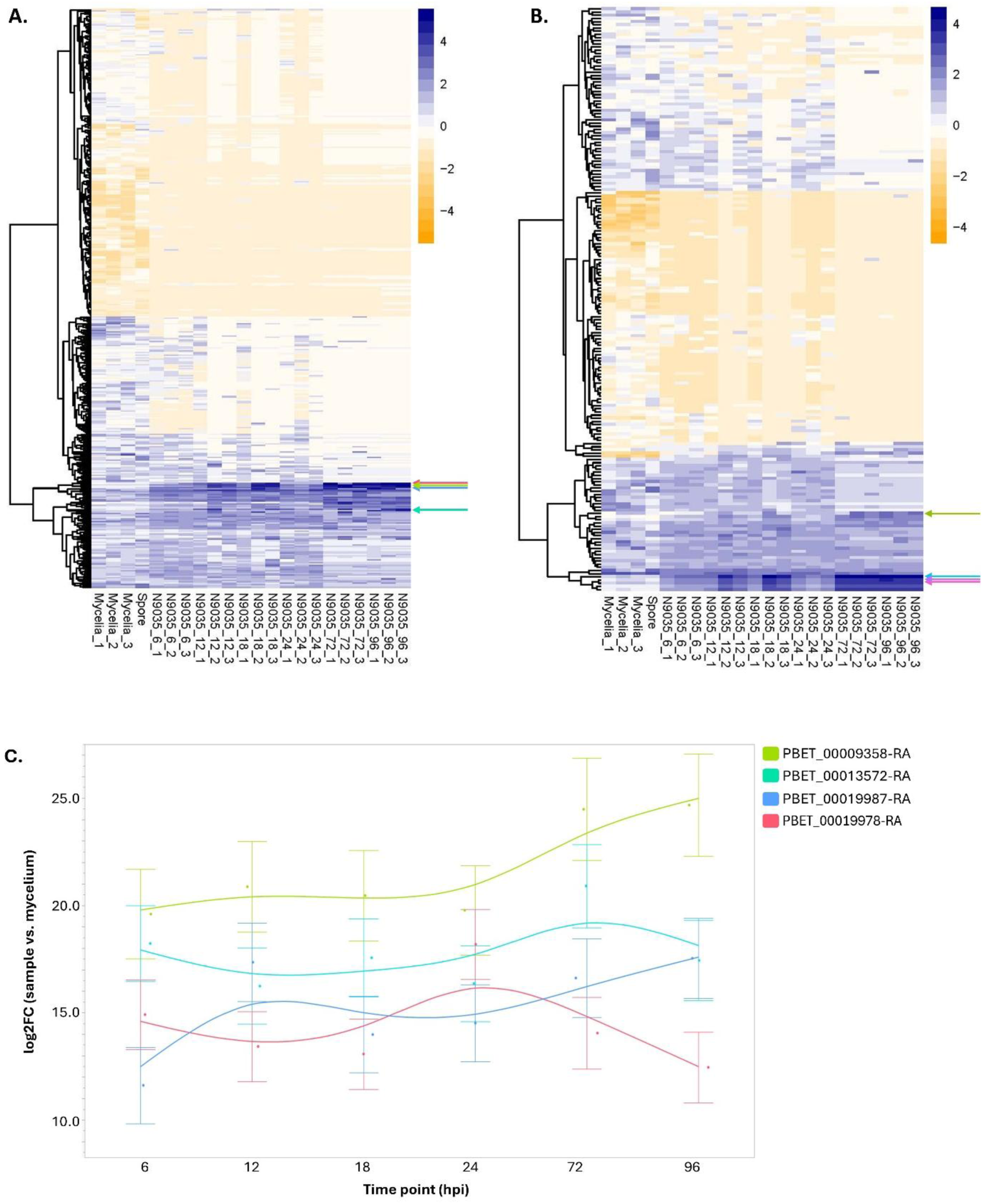
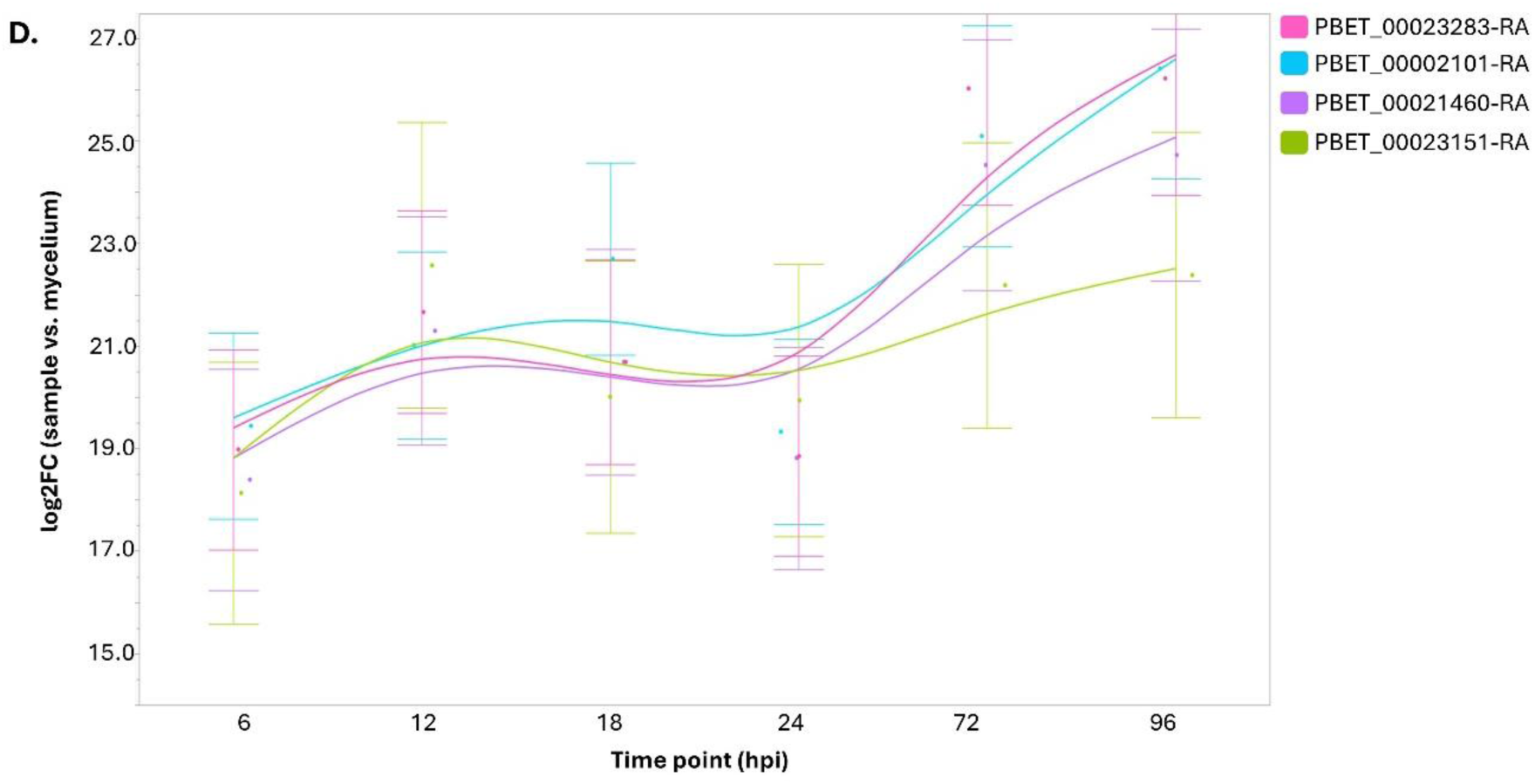
Expression patterns of *P. betacei* effectors across its infection cycle in *S. betaceum*. Heatmap of the mean-centered z-scores of *P. betacei* **(A)** RxLR effectors and **(B)** CRN effectors. Colored arrows indicate the genes shown in figures C and D. Expression patterns of four members from the RxLR group of effectors **(C)** and the CRN group of effectors **(D).** Dots represent the mean values of expression (log2fold change values) from RNAseq, for each gene at each time point, when compared to their expression in the mycelium. A trend line is projected over the data points and bars represent the standard error (SE) biological replicates.

## Discussion

In this study, we used an integrative approach that leveraged epidemiological, cellular and molecular approaches to describe the infection cycle of *P. betacei* on tree tomato. The data collected highlighted that *P. betacei* strains display notable differences when infecting a susceptible host, while *P. infestans* strains show a more stable isolate distribution. Furthermore, a detailed investigation of the infection cycle of *P. betacei* and *P. infestans* allowed us to clearly identify the biotrophic and necrotrophic stages at the cellular level, by visualizing infected material, and at the molecular level, by assessing the expression dynamics of two marker genes. Even so, we identified a changing transcriptomic landscape of *P. betacei* when in its biotrophic versus its necrotrophic stage, *P. betacei* exerts a hemibiotrophic infection on tree tomato characterized by a long biotrophic period, the presence of feeding structures, and a late necrotrophic stage where the pathogen sporulates to start a new cycle.

All *P. betacei* isolates tested in this study infected and caused disease on a susceptible host; however, we observed a large variation in their phenotypes. Such a level of phenotypic variation is most probably a consequence of the tree tomato not yet being fully domesticated (Acosta-Quezada et al. 2011). Considering that *P. betacei* has been described as a specialist of this host (Mideros et al. 2018) and that the lack of domestication would mean that there is no established breeding program for it (Acosta-Quezada et al. 2011), broad genetic diversity of the host would be expected. Therefore, the thriving diversity of *S. betaceum* could be driving increased variability in *P. betacei* as well. This could impact how the disease spreads in the field and how it can be managed in such a diverse environment.

Additionally, previous studies have reported variability in aggressiveness among isolates belonging to the same species or sharing similar genotypes (Pariaud et al. 2009). For example, a previous study demonstrated that significant variation for foliar aggressiveness (measured as lesion expansion rate, latent period, sporulation and infection efficiency) exists within the Northern Ireland population of *P. infestans* when inoculated on the same potato cultivar (Carlisle et al. 2002). In that case, the variation was explained in terms of the geographic origin of each strain. Similar findings were reported for the leaf rust pathogen *Puccinia striiformis f. sp. tritici* (Milus, Seyran and Mcnew 2006) where isolates belonging to the same pathotype exhibited different latency periods. This strong reliance on diversity may be interpreted as the result of balancing selection on epidemiological components that may be involved in trade-offs at different stages of the life cycle. In other words, as a way of diminishing the risk of extinction in unstable environments by having different ways to respond (Andrivon et al. 2013; Chakraborty 2013). An example of an unstable environment would be the domestication process that *S. betaceum* has been going through. Additionally, we can attribute this variability to the performance of our experiments on tree tomato plants given that the accession used in this study has not been accurately phenotyped or genotyped, and thus we relied on undocumented susceptibility information that can bias our results.

A key aspect for effective control measures of *Phytophthora* diseases is the comprehensive understanding of the infection at cellular and molecular levels (Hardham 2001). Using light and scanning electron microscopy, coupled with gene expression analyses, we determined that *P. betacei* follows a hemibiotrophic infection cycle. It first attaches firmly to the surface, then penetrates the tissue, and then colonizes it while taking up nutrients for growth and sporulation. However, we found that it has a long biotrophic phase until around 24 hpi. This is comparable with observations made in *P. infestans, P. capsici, P. palmivora, and P. sojae* in their natural hosts (Ali et al. 2017; Evangelisti et al. 2017; Jupe et al. 2013; Kunjeti et al. 2012; Ye et al. 2011; Zuluaga et al. 2016). For example, *P. infestans* develops mature haustoria 12hpi and invades a susceptible potato host 72 hpi, when the production of new asexual structures (sporangia) is evident (Grenville-Briggs and van West, 2005). However, biotrophy was much shorter for *P. infestans* on tree tomato, occurring only for 24 hours. An extended biotrophy could be an indicator of how adapted are different isolates to their hosts (Kröner et al. 2017) and could suggest the presence of specific adaptations required by *P. betacei* to thrive on tree tomato.

The use of molecular markers contributed to a better understanding of the colonization process and validated the phenotypic observations. On the one hand, the presence of biotrophic feeding structures (haustoria) was accompanied by the upregulation of the gene encoding the haustorium membrane protein (*Hmp1*). *Hmp1* has been proposed to impact protein stability of the pathogen’s membrane and localizes only in regions where infection vesicles and haustoria are observed (Avrova et al. 2008). These observations support *P. betacei* and *P. infestans* having a compatible interaction with tree tomato. Usually, the degree of compatibility of the interaction even influences the number of haustoria that are formed (Hardham 2001). Moreover, the development of haustoria has been associated with upregulation of *Hmp1* in *P. infestans, P. palmivora* and *P. capsici* (Ah Fong and Judelson 2003; Jupe et al. 2013; Le Fevre et al. 2016). Comparably, the expression profile of cell cycle phosphatase 14 (*Cdc14)*, accurately supported the results of a later necrotrophic phase by showing to be upregulated at 96 hpi for *P. betacei* and 72 hpi for *P. infestans*. This protein plays an essential role during asexual sporulation and biogenesis of the flagellar apparatus (Ah-fong and Judelson 2011; Ah Fong and Judelson 2003). Given that the formation of sporangia is essential for the propagation of *Phytophthora,* this is usually associated with necrotrophy. *Cdc14*, has also been found to be upregulated later in the infection cycle and during sporangial production in *P. infestans, P. palmivora* and *P. capsici* (Ah Fong and Judelson 2003; Jupe et al. 2013; Le Fevre et al. 2016).

RNAseq data provided evidence of a changing transcriptional landscape for *P. betacei* as the infection progresses, just as has been described for *P. infestans*, *P. sojae*, or *P. cinnamomi* across their life cycles *in planta* (Fong, Kim and Judelson 2017; Lin et al. 2014; Reitmann, Berger and van den Berg 2017). This supports the hypothesis that the infection features stage-specific transcriptional programs (Randall et al. 2005). Additionally, it provides an important complement to the transcriptional landscape of *S. betaceum* throughout the infection of *P. betacei* (Bautista et al. 2021). In that study, most of the DEGs were involved in plant defense and changed along the cycle in parallel to the changes occurring in the pathogen. Lastly, genes that are important for infection progression in other *Phytophthora* species were identified as relevant for *P. betacei* as well.

For example, the glucanase inhibitor protein (GIP) is known to inhibit the activity of endoglucanases involved in the plant defense against infection of *P. cinnamomi* (Martins et al. 2014). Therefore, its slight upregulation at the beginning of the cycle may represent an early effort of *P. betacei* to counteract plant defense. As expected, this was then followed by upregulation again in the necrotrophic stage, when the plant has established most of its response to the pathogen. Similarly, the suppressor of necrosis 1 (*SNE1*), a widely known RXLR effector of *Phytophthora* involved in managing the hemibiotrophic transition (Kelley et al. 2010) was also identified as differentially expressed along the infection cycle of *P. betacei*. This protein is usually expressed during the biotrophic stage given that it inhibits cell death (Zuluaga et al. 2016). In this study, it was upregulated early in the cycle, as expected, but then increased its expression again towards 24 hpi and onwards. This may be explained by the elongated hemibiotrophic transition described for *P. betacei* but it would be necessary to continue monitoring its expression later in the cycle to determine its pattern in later stages.

Lastly, monitoring the expression profiles of *P. betacei* effectors along the cycle allowed us to further confirm that this pathogen has an elongated hemibiotrophic transition when compared to *P. infestans.* Since RxLR effectors are known for their modulation of and impact on host defenses (Zuluaga et al. 2016), they are known to have variable patterns of expression across the cycle since they respond to how the interaction between pathogen and host progresses (Birch et al. 2009). Therefore, the multiple expression patterns observed for RxLRs in this study are supported by their natural variability, but also reinforce that the time points of 72 and 96 hpi are certainly a necrotophic stage for *P. betacei* on *S. betaceum*. In fact, CRN effectors known to generate a crinkling phenotype in the leaves and to drive necrosis (Stam et al. 2013) tended to be upregulated at 72 hpi and further. Overall, our results support a disease model where *P. betacei* behaves as a biotroph between 0 and 18hpi and starts its hemibiotrophic transition that lasts until before 72 hpi, when the infection is clearly necrotic.

This study highlights the importance of using the combined effect of several epidemiological parameters in the characterization and selection of *P. betacei* strains, as greater differences can be detected when the contribution of each variable is leveraged (Miller, Johnson and Hamm 1998). Natural variation in the infection phenotype has been observed within the *Phytophthora* genus and might be an indicator of the pathogen’s adaptation level. Furthermore, a clear hemibiotrophic infection cycle was confirmed by the presence of haustoria during an asymptomatic period, followed by necrosis and sporulation. The mechanisms behind the extended biotrophic stage remain unexplored and require special attention as they might be involved in the host specificity of *P. betacei*. Our data on the transcriptional landscape at each point of the infection can certainly support this understanding. However, it also highlighted the importance of generating high-quality genetic resources for *S. betaceum* and *P. betacei*, given that there is no available genome of *S. betaceum*. Overall, this study serves as a bridge to continue deciphering the processes this *Phytophthora* shares with others while highlighting its particularities.

## ACKNOWLEDGMENTS

The authors would like to thank Anna Gogleva for the invaluable help regarding the transcriptional analyses. This work was supported by The Royal Society International Exchanges Award 2016/R2 Ref. IE160666.

## SUPPLEMENTARY TABLES

**Supplementary Table 1.** Median value of each evaluated epidemiological parameter among strains. Ten *P. betacei* and three *P. infestans* strains were inoculated on susceptible tree tomato (Comun accession) plants. The first and third quartiles are indicated in parentheses as a dispersion measurement. The size of the sample (n) might differ among strains due to the lack of symptoms and signs in some of the plants.

| Epidemiologica<br>l parameter | Species |  |  |  |  |  |  |  |  |  |  |  |  |
| --- | --- | --- | --- | --- | --- | --- | --- | --- | --- | --- | --- | --- | --- |
|  | *P. betacei* |  |  |  |  |  |  |  |  |  | *P. infestans* |  |  |
|  | P9127<br>n =9 | P8084<br>n=5 | P8077<br>n =9 | P9151<br>n =9 | P9153<br>n =9 | P8029<br>n =6 | N9035<br>n =9 | N9070<br>n =8 | N9022<br>n =6 | SBC3#<br>7<br>n =8 | Z-32<br>n =9 | VPC7#<br>10<br>n =6 | STT23<br>5<br>n =8 |
| **Lesion Area**<br>**(mm<sup>2</sup>)** | 33.64<br>(16.2-42.36) | 8.16<br>(5.28-9.130) | 20.36<br>(16.67-21.56) | 18.77<br>(15.24-54.28) | 60.42<br>(49.41-73.75) | 18.50<br>(4.70-35.75) | 99.05<br>(17.0 –<br>114.50) | 18.80<br>(18.0-19.0) | 6.68<br>(3.11-10.9) | 13.14<br>(9.621-17.8) | 64.45<br>(17.21-72.97) | 51.63<br>(45.7-101) | 54.33<br>(25.3-75.4) |
| **Lesion Growth**<br>**Rate (mm<sup>2</sup>h<sup>-1</sup>)** | 0.15<br>(0.07-0.19) | 0.037<br>(0.02-0.06) | 0.094<br>(0.07-0.09) | 0.087<br>(0.07-0.25) | 0.28<br>(0.22-0.34) | 0.085<br>(0.02-0.16) | 0.45<br>(0.07-0.53) | 0.087<br>(0.08-0.09) | 0.031<br>(0.014<br>- 0.05) | 0.061<br>(0.04-0.08) | 0.298<br>(0.07-0.33) | 0.239<br>(0.21-0.46) | 0.251<br>(0.11-0.35) |
| **Incubation**<br>**Period (hpi)** | 96.00<br>(53.3-120) | 122<br>(100-141) | 36<br>(34.29-48) | 48<br>(48.0-48.0) | 58.66<br>(55.2-64.0) | 120<br>(114-120) | 104<br>(60.0-112) | 90<br>(71.5-106) | 117<br>(99.21-129) | 75.27<br>(69.0-93.0) | 48<br>(45.0-72.0) | 24<br>(24.0-40.0) | 46.50<br>(42.0-49.5) |
| **Sporulation Capacity (sp mm<sup>-2</sup>)** | 509.69<br>(405.1-815.7) | 308.45<br>(0-730.8) | 412.00<br>(201.1-412.0) | 201.14<br>(0-1896.0) | 503.45<br>(354.3-506.6) | 0<br>(0-110.5) | 476.73<br>(417.5-736.1) | 0<br>(0-264.5) | 238.36<br>(0-827.8) | 82.75<br>(0-1388) | 24.55<br>(0-39.53) | 47.14<br>(47.17-69.53) | 287.1<br>(0-69.12) |
| **Latency Period (hpi)** | 192<br>(162-194) | 108<br>(0-216) | 216<br>(185.1-216) | 162<br>(0-216) | 216<br>216 | 0<br>(0-81) | 168<br>(144-168) | 0<br>(0-216) | 108<br>(0-216) | 99<br>(0-198) | 216<br>(0-216) | 144<br>(0-154.3) | 108<br>(0-216) |
| **Sporulation Rate (sp hpi<sup>-1</sup>)** | 39.78<br>(34.72-119.6) | 19.89<br>(0-426.6) | 46.29<br>(17.36-46.30) | 54.65<br>(0-262.1) | 78.39<br>(67.52-114.1) | 0<br>(0-781.2) | 61.73<br>(31.83-3981) | 0<br>(0-18.33) | 15.91<br>(0-32.07) | 32.55<br>(0-165.5) | 5.06<br>(0-7.23) | 11.57<br>(0-25.63) | 12.86<br>(0-40.9) |

**Supplementary Table 2.** ***Phytophthora betacei* and *Phytophthora infestans* strains used in this study.**

| Strain ID | Host | Species | Location | Reference |
| --- | --- | --- | --- | --- |
| **P9127** | *Solanum betaceum* | *P.betacei* | Putumayo, Colombia | (Mideros et al. 2018) |
| **P8084** | *S. betaceum* | *P.betacei* | Putumayo, Colombia | (Mideros et al. 2018) |
| **P8077** | *S. betaceum* | *P.betacei* | Putumayo, Colombia | (Mideros et al. 2018) |
| **P9151** | *S. betaceum* | *P.betacei* | Putumayo, Colombia | (Mideros et al. 2018) |
| **P9153** | *S. betaceum* | *P.betacei* | Putumayo, Colombia | (Mideros et al. 2018) |
| **P8029** | *S. betaceum* | *P.betacei* | Nariño, Colombia | (Mideros et al. 2018) |
| **N9022** | *S. betaceum* | *P.betacei* | Nariño, Colombia | (Mideros et al. 2018) |
| **N9035** | *S. betaceum* | *P.betacei* | Nariño, Colombia | (Mideros et al. 2018) |
| **N9070** | *S. betaceum* | *P.betacei* | Nariño, Colombia | (Mideros et al. 2018) |
| **Sbc3#7** | *S. betaceum* | *P.betacei* | Cundinamarca,<br>Colombia | (Chaves et al. 2016) |
| **VCP7#10** | *S. tuberosum* | *P.infestans* | Cundinamarca,<br>Colombia | (Chaves et al. 2016) |
| **STT235** | *S. tuberosum* | *P.infestans* | Nariño, Colombia | (Parra et al. 2016) |
| **Z-32** | *S. phureja* | *P.infestans* | Zipaquirá, Colombia | (Cárdenas et al. 2012) |

**Supplementary Table 3.**
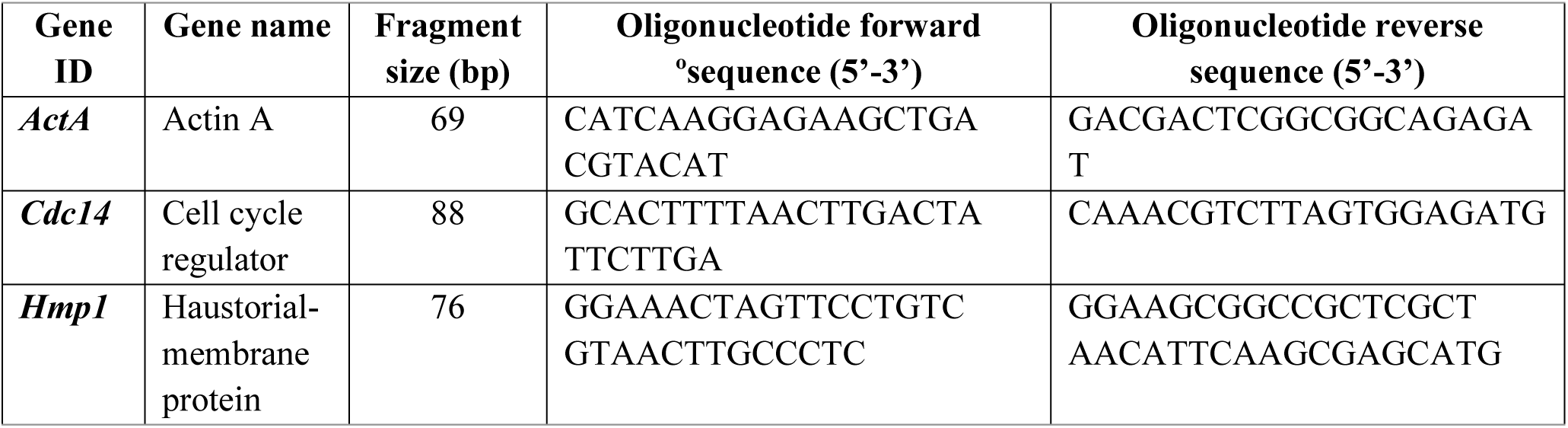
Sequence of specific qRT-PCR oligonucleotides used to assess marker gene expression during the infection cycle of *P. betacei* and *P. infestans* on tree tomato.

**Supplementary Table 4.**
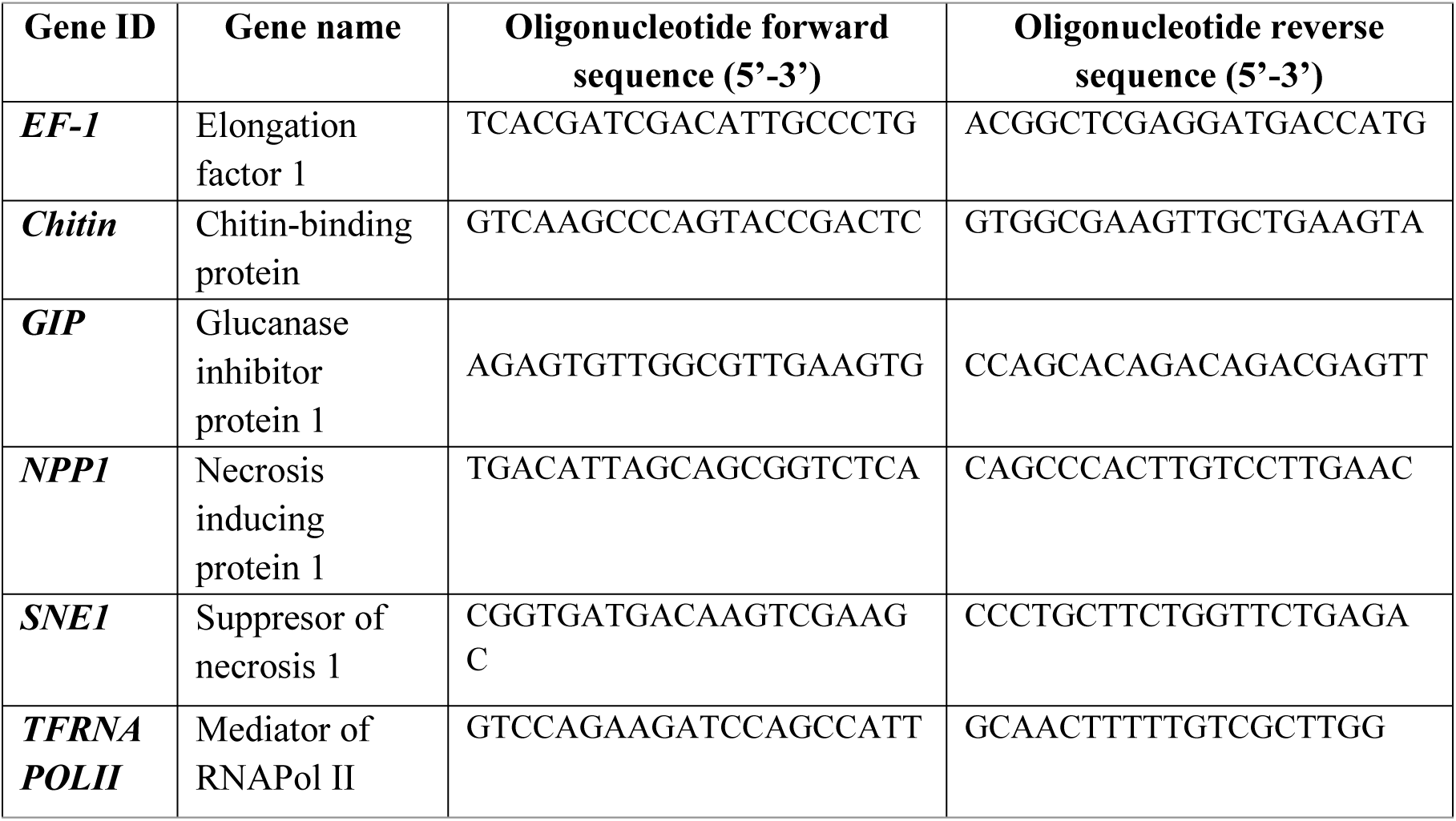
Sequence of specific qRT-PCR oligonucleotides used to validate the gene expression detected via RNAseq during the infection cycle of *P. betacei* on tree tomato.

